# A native mass spectrometry-based assay for rapid assessment of the empty/full capsid ratio in AAV gene therapy products

**DOI:** 10.1101/2021.07.06.451264

**Authors:** Lisa Strasser, Tomos E. Morgan, Felipe Guapo, Florian Füssl, Daniel Forsey, Ian Anderson, Jonathan Bones

## Abstract

Adeno-associated virus (AAV)-based cell and gene therapy is a rapidly developing field, requiring analytical methods for detailed product characterization. One important quality attribute of AAV products that requires monitoring is the amounts of residual empty capsids following downstream processing. Traditionally, empty and full particles are quantified via analytical ultracentrifugation as well as anion exchange chromatography using ultraviolet or fluorescence detection. Here, we present a native mass spectrometry-based approach to assess the ratio of empty to full AAV-capsids without the need for excessive sample preparation. We report rapid determination of the amount of empty particles in AAV5 and AAV8 samples, with results correlating well with more conventional analysis strategies, demonstrating the potential of state-of-the-art mass spectrometry for the characterization of viral particles.

---

Adeno-associated virus (AAV)-based cell and gene therapy is evolving rapidly. Since the first AAV-based product was approved by the European Medicine Agency (EMA) in 2012, more than 150 AAV related clinical trials were listed on ClinicalTrials.gov^1^. However, despite this impressive progress, analytical methods to monitor quality attributes of recombinant AAV (rAAV)-based products have not advanced with the same speed in recent years.

AAVs are composed of a protein capsid encapsulating a ∼4.7 kb single-stranded DNA genome. The capsid is assembled by 60 copies of the viral proteins VP1, VP2, and VP3 in a ratio of approximately 1:1:10, building a capsid of ∼3.8 MDa^2^. Of particular concern during the production of rAAV is the amount of empty capsids present which is not only important for administering the correct dosage but also to account for concerns regarding potential unwanted immune response caused by empty capsids^3^. There are various methods available to quantify empty and full capsids^4,5^, the most common being analytical ultracentrifugation (AUC)^6^ as well as, more recently introduced, anion-exchange chromatography (AEX)^7,8^. While these tools have been shown to successfully separate empty and full capsids of various serotypes, absorbance-based methods still face certain limitations. Even though UV-detection at 260 and 280 nm can be used to differentiate empty and full capsids during AEX separation, it is known to lack the required sensitivity which is of key importance when working with AAV-samples of low concentration. Furthermore, a response factor is needed for correction during quantitation using UV-absorbance. This can be avoided by using fluorescence detection which, however, does not allow for an unambiguous identification of empty and full capsids^6,9,10^. This problem could potentially be circumvented using a mass spectrometry (MS)-based approach.

In recent years, the application of mass spectrometry for the analysis of AAV particles has gained tremendous interest^11,12^. Using intact native MS analysis allowed for the determination of the molecular mass of AAV capsids and thereby also revealed the enormous inherent heterogeneity of viral particles^13^. Still, this heterogeneity in combination with the high molecular weight of intact AAV capsids poses significant analytical challenges. Conventional non-isotopically resolving MS requires detection and resolution of multiple consecutive charge states for deconvolution and is therefore only applicable to samples of limited complexity. While the use of native conditions results in less charges and higher spatial resolution in the m/z space^14^, it is currently still not possible to gain charge state resolution for intact AAV particles.

Even though not yet fully commercially available, charge detection mass spectrometry (CDMS) addresses this problem by directly measuring the mass of individual ions, enabling analysis of high molecular weight species at a level that has not previously been possible^15-18^. Interestingly, CDMS analysis has shown that empty and full AAV particles have a similar charge state distribution yet differ significantly in their mass^16,19^.

Here, we exploit this information to determine the empty/full ratio of rAAVs using conventional MS under native conditions. Observed signal clusters derived from empty and full AAV capsids were assigned and facilitated area-based quantitation, resulting in an easy-to-implement assay utilizing standard instrumentation readily available in many characterisation labs.

## RESULTS AND DISCUSSION

Empty and full AAV5 were analysed by native direct infusion mass spectrometry, resulting in signal clusters in a range of *m/z* 18,000-23,000 and 23,000-32,500, respectively (Figure 1, a and b). Importantly, empty reference material might contain full capsids and vice versa, resulting in additional signal cluster as can be seen especially in Figure 1a (m/z > 23,000). Still, full and empty capsids appeared to follow the trend previously obtained by CDMS and appeared in different m/z regions, indicating the same charge while being of different mass^16,19^. Assuming an average charge of +150 to +160, as was reported previously^19^, the mass of AAV5 empty was found to be between 2.9 and 3.1 MDa while full capsids appeared to have a mass of 3.8–4.1 MDa. Thus, the observed mass difference between full and empty particles correlates well with the mass of the incorporated cargo genome (2.5 kb, approx. 800 kDa). Noteworthy, full capsids appeared in a broader cluster, indicating higher heterogeneity due to the incorporated ssDNA.

**Figure 1.**
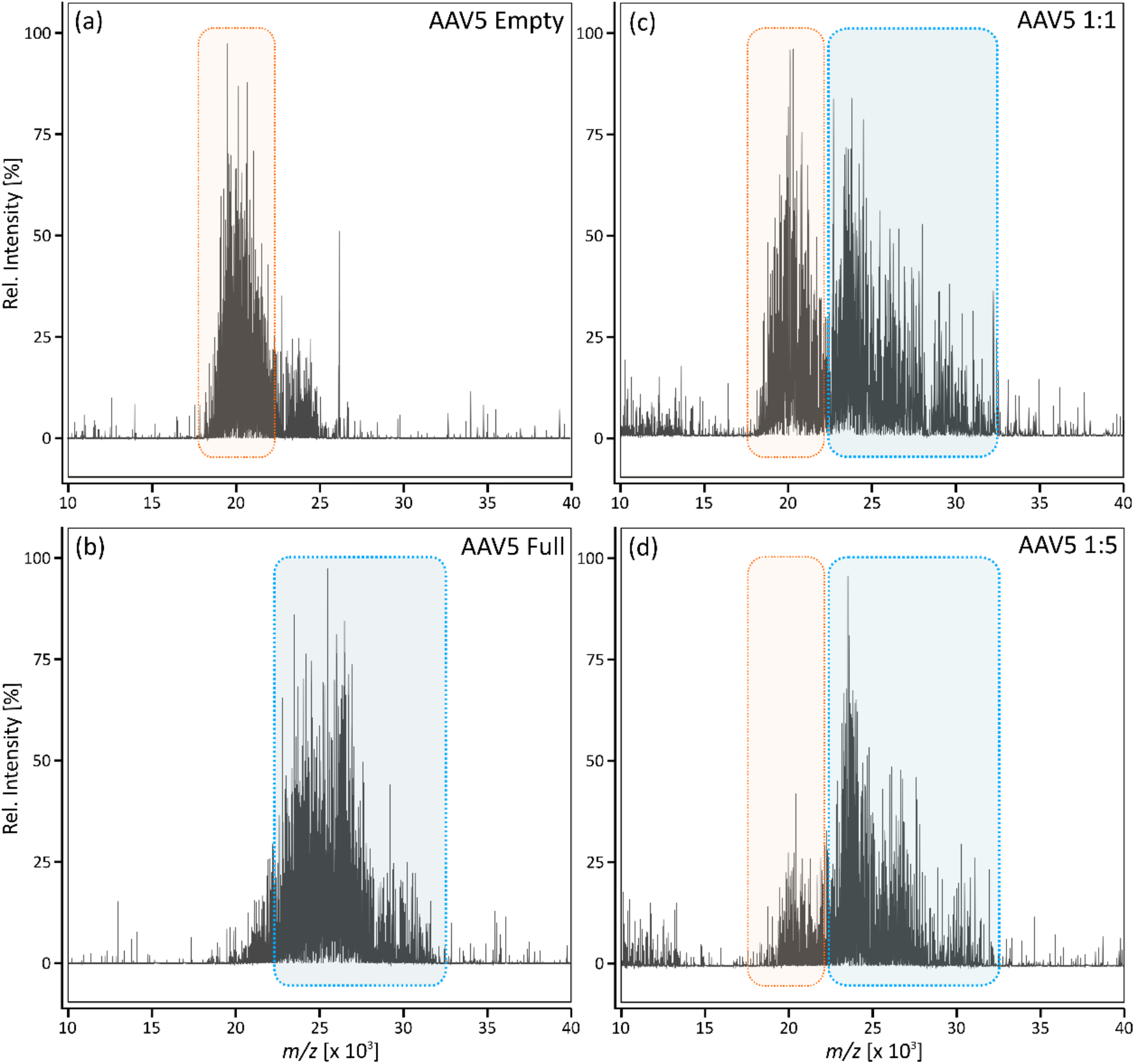
Native direct infusion MS of AAV5. Empty (a) and full (b) AAV5 reference material was analysed individually as well as in a volumetric mixture of 1:1 (c) and 1:5 (d). Shown are averaged spectra after 5 minutes of data acquisition. Signal cluster derived from empty AAV is highlighted in orange, full is highlighted in blue.

Next, mixtures of full and empty AAV5 were analysed. As shown in Figure 1c and d, observed signal clusters still appeared in the same *m/z* region while the relative abundance changed, correlating to the respective mixture of 1:1 and 1:5 (*V/V*).

Interestingly, spray stability during static nanoESI infusion was observed to differ considerably depending on the AAV serotype with AAV5 being particularly difficult to analyse over extended periods of time. Whether this is due to sample stability in ammonium acetate requires further investigations. Nevertheless, while spray stability is a crucial factor during CDMS analysis where extensive data acquisition times are required, it did not noticeably affect the quality of the presented data, as acquisition times were merely 5 minutes per measurement. To the best of our knowledge, this is the first time, spectra obtained from empty as well as full AAV5 are reported.

To test the method for applicability with different serotypes, the same analysis was performed for AAV8. Figure 2 is showing the results obtained for a 1:1 and 1:5 mixture of empty and full capsids of AAV8. Acquired charge on ionization caused shifts in the *m/z* distribution dependent on serotype. Therefore, AAV8 generally appeared at a higher *m/z* range but intensities corresponding to full and empty particles still changed according to their concentration.

**Figure 2.**
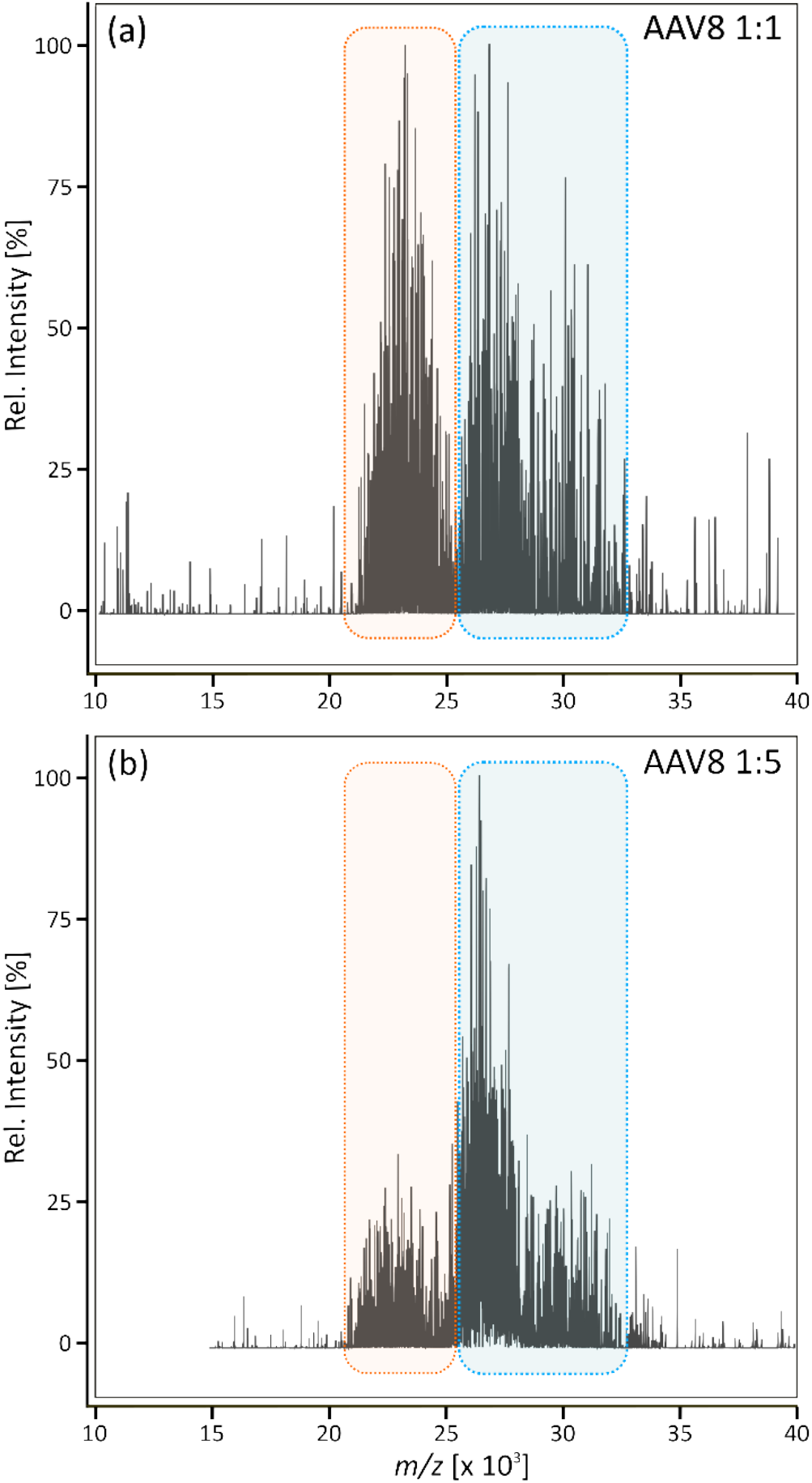
Intact native MS analysis of AAV8. Full and empty reference material was mixed in a ratio of 1:1 (a) and 1:5 (b). Signal cluster derived from empty AAV is highlighted in orange, full AAV is highlighted in blue.

Importantly, the analyses performed did not result in charge state resolution and do therefore not allow for direct determination of the accurate mass of AAV5 and AAV8. However, differential signal clusters were clear and allowed for the relative quantification of full and empty capsids. To determine the empty to full ratio of the analysed samples, corresponding cluster areas were measured using ImageJ. Resulting data was exported for further analysis, the results obtained are shown in Table 1.

**Table 1.** Empty to full ratio assessment of AAV5 and AAV8. AAV reference material was mixed in a ratio of 1:1 and 1:5, respectively. Samples were analysed via native MS and resulting data was analysed using ImageJ. Signal clusters corresponding to empty and full capsids were measured and subsequently used to calculate the ratio of empty to full as well as the percentage of full capsids. Additionally, empty and full capsids were separated using AEX and peak areas were used to calculate the relative amount of full AAV in %.

| Sample | Empty/Full (V/V) | Ratio | % Full (MS) | % Full (AEX) |
| --- | --- | --- | --- | --- |
| AAV5 | 1:1 | 1.56 | 61.02% | 62.66% |
| AAV5 | 1:5 | 4.60 | 82.16% | 77.54% |
| AAV8 | 1:1 | 1.41 | 58.50% | 58.21% |
| AAV8 | 1:5 | 3.28 | 76.62% | 73.94% |

As indicated in Table 1, a volumetric mixture of empty and full reference material in a ratio of 1:1 was found to contain approximately 60% full capsids while a mixture of 1:5 contained about 82% full capsids which correlates well with what is to be expected. Furthermore, the obtained results agree with numbers obtained by fluorescence detection using AEX separation (below 5% variation, data shown in supplementary Figure 1). Reproducibility of native MS analysis was evaluated by triplicate analysis of AAV5 samples (Figure S2 and Table S1 in the supplement). Thereby, the standard deviation was found to be below 1.05% indicating high consistency. This clearly demonstrates that conventional MS under native conditions can be used to reliably assess the empty to full ratio of AAV samples. Further modifications of the presented method such as e.g. the use of charge reduction to increase spatial resolution might also enable quantitation of partially filled capsids. In any case, while the required analysis time is significantly reduced compared to standard AEX, MS analysis furthermore allowed for an unambiguous assignment of signals to full and empty capsids without the need for further experiments as well as for the reliable evaluation of their relative abundances.

Finally, to demonstrate applicability of the presented method for samples derived from downstream processing, an in-process sample of AAV5 was analysed (Figure 3). Thereby, 53.45% of the capsids were found to be full resulting in an empty/full ratio of 1.15. Results correlate with the amount of full capsids determined via AUC, carried out by Pharmaron (data not shown).

**Figure 3.**
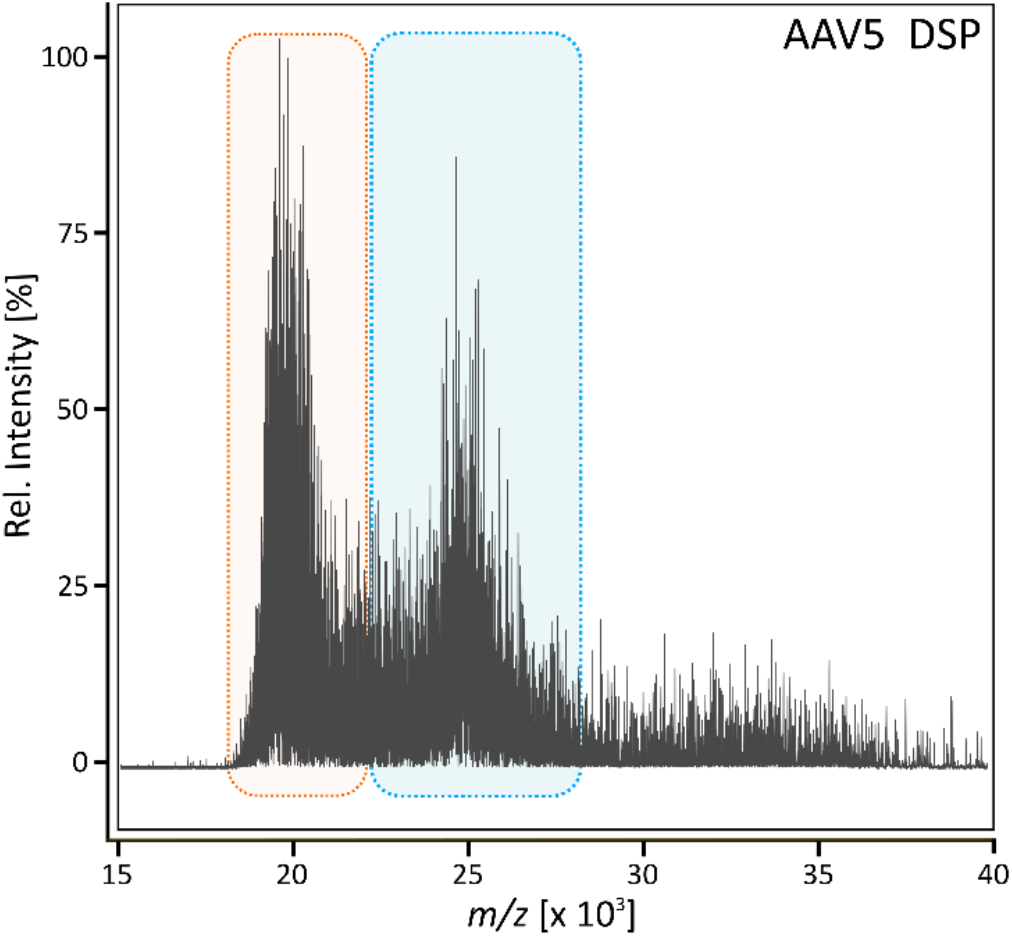
Intact native MS analysis of AAV5 derived from down-stream processing (DSP). Signal cluster derived from empty AAV5 is highlighted in orange, full AAV is highlighted in blue.

## CONCLUSION

Direct infusion native mass spectrometry was used to measure the relative abundance of AAV5 and AAV8 capsids with and without cargo DNA. Signal clusters derived from empty and full capsids were clearly differentiated and their relative abundances correlated well with expected values based on the deliberate generation of samples with varying empty/full ratio. Importantly, the approach presented has shown applicability for multiple AAV serotypes as well as samples derived from downstream processing. Taken together, the results presented clearly demonstrate the potential of using commercially available state-of-the-art mass spectrometry for the analysis of crucial quality attributes of high molecular weight analytes, such as AAVs. Although, due to sample size and complexity, AAV charge states remained unresolved, *m/z* spacing of the filled and unfilled capsid allowed for relative quantitation, offering a great prospect for the coupling of MS analysis strategies to chromatographic separation techniques. Thereby, deeper and more accurate analysis through coupling of analytical techniques or further charge detection MS (CDMS) methods will allow more accurate mass determination of viral capsids. In any case, further technological developments will fully enable detailed characterization of next generation biotherapeutics such as AAV.

## ASSOCIATED CONTENT

### Supporting Information

The Supporting Information is available free of charge on the ACS Publications website.

Additional experimental details, materials, and methods, about native MS analysis and anion exchange chromatography (AEX). Results of empty/full separation using AEX. Replicate analysis of AAV5 using native MS analysis.

## AUTHOR INFORMATION

### Author Contributions

The manuscript was written through contributions of all authors. / All authors have given approval to the final version of the manuscript.

### Notes

The authors declare no competing financial interest.

## ACKNOWLEDGMENT

The authors greatly acknowledge funding from Enterprise Ireland under the Innovation Partnership Program IP/2018/0753 and support from Abbvie and Pharmaron.

## Table of Contents artwork

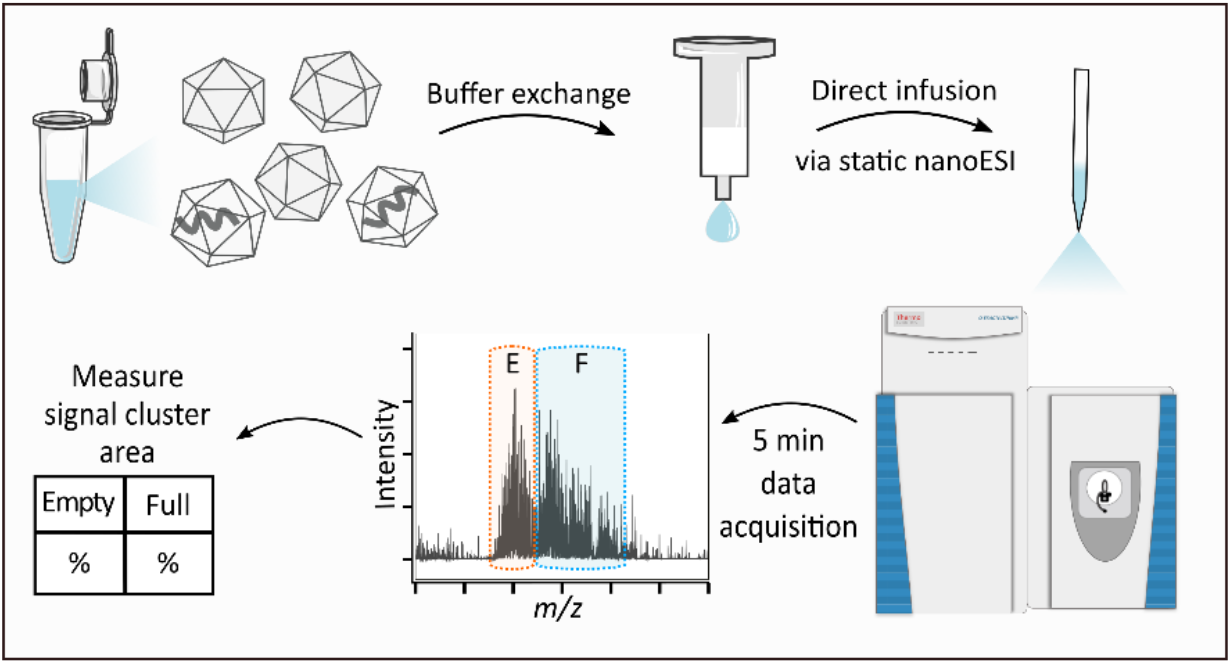

## Supporting Information

### 1. EXPERIMENTAL SECTION

AAV5 and AAV8 full and empty references were purchased from Virovek (Hayward, CA, USA). Both serotypes are derived from Sf9 insect cells using a baculovirus expression system. Ammonium acetate (99.999% trace metals basis), bis-tris propane (BTP) and trimethylammonium chloride (TMA) were purchased from Sigma Aldrich (Wicklow, Ireland) and LC-MS grade Water was obtained from Fisher (Dublin, Ireland).

#### 1.1. Native MS analysis

Prior to analysis, AAV samples were buffer exchanged into aqueous ammonium acetate (100 mM, pH 6.8-7.0) using Bio-Spin® P-6 Gel Columns (Bio-Rad Laboratories, Hercules, CA, USA) according to the manufacturer’s instructions. The final sample concentration was between 5 × 10^12^ – 1 × 10^13^ viral particles/mL, an aliquot of 5 µL was loaded into double-coated borosilicate nESI emitter tips (Thermo Scientific, Hemel Hempstead, UK). Samples were analysed on a Q Exactive UHMR hybrid quadrupole-Orbitrap mass spectrometer equipped with a Nanospray Flex ion source (Thermo Fisher Scientific, Bremen, Germany). Instrument parameters were optimized using a sample with known concentration of empty and full capsids to avoid biased results. Data was acquired in positive ion mode using a resolution setting of 25,000 (at *m/z* 200), 10 microscans and an automatic gain control of 1e6 with a maximum injection time of 200 ms. Spray voltage was 1.5 kV, capillary temperature was set to 250°C and the S-lens RF level was 200. The ion transfer target *m/z* and detector optimization were set to “high *m/z*”. In-source trapping was enabled with a desolvation voltage of – 100 V and a source DC offset of – 50 V. Extended trapping of particles was carried out in the HCD cell with a HCD energy of 150 V. Sulphur hexafluoride (SF6) was used as collision gas at 4.0 e-10 mbar (UHV readout). Data was acquired for 5 min with transient averaging enabled.

#### 1.2. Anion exchange chromatography

Empty and full separation of AAV samples was conducted on a ProPac SAX-10 1 × 50 mm column (Thermo Fisher Scientific, Sunnyvale, CA, USA) using fluorescence detection (Ex280/Em340) on a Vanquish Horizon UHPLC system (Thermo Scientific, Germering, Germany). Buffer A was 20 mM BTP at pH 9.0 and buffer B was 20 mM BTP with 1 M TMA at pH 9.0. At a flow rate of 0.15 mL/min and a temperature of 30°C, empty and full capsids were separated using the following conditions: samples were loaded onto the column at 0.1% B followed by an isocratic hold for 3 min. Separation was done using a linear gradient of 1-50% B in 15 min followed by a wash step at 90% B for 1.5 min and column re-equilibration at 0.1% B for 20 min.

## 2. RESULTS

**Supplementary Figure 1.**
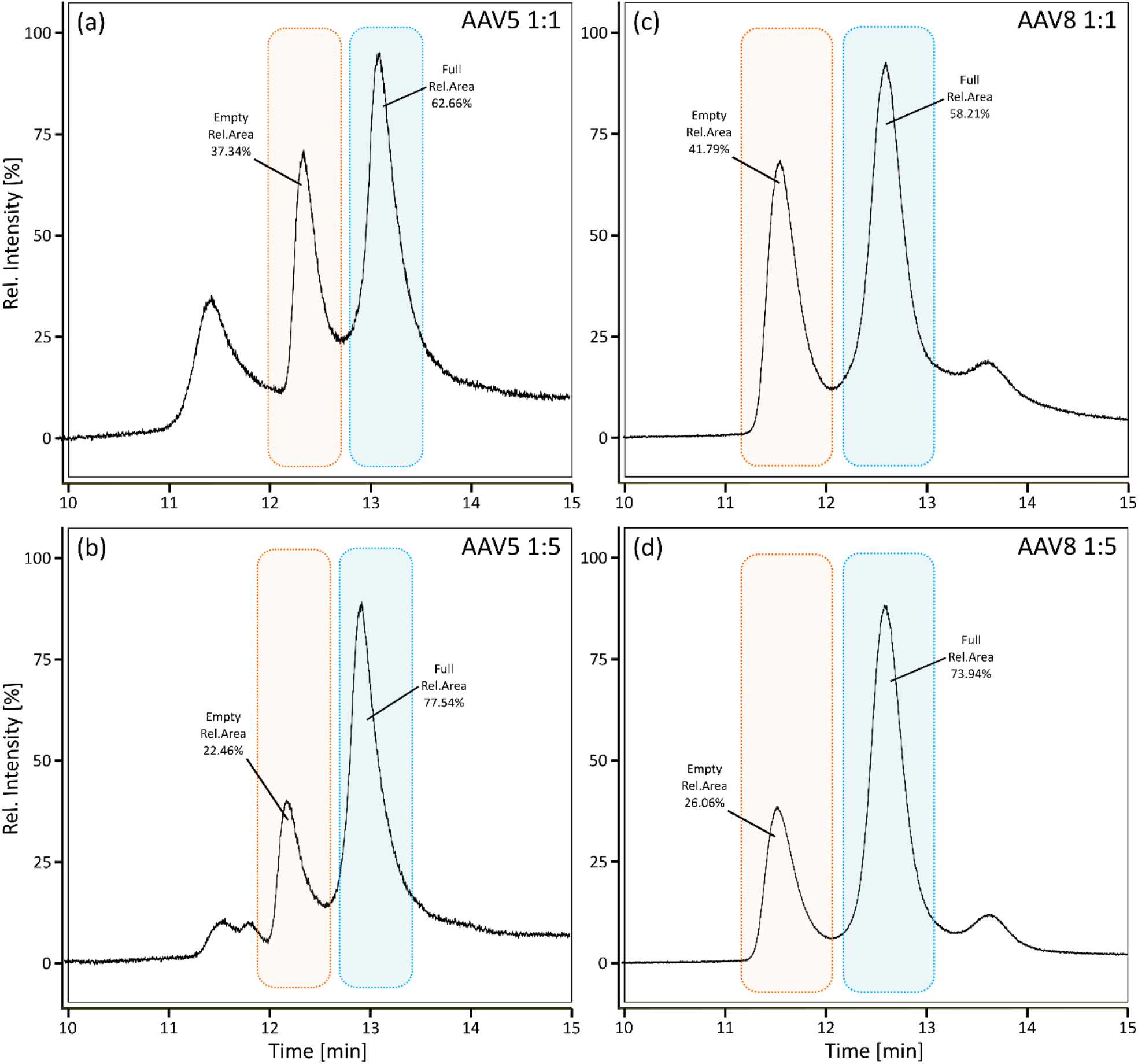
Anion exchange chromatography-based LC-separation of empty and full AAV capsids. AAV5 reference material was analysed in a volumetric mixture of 1:1 (a) as well as 1:5 (b). Similar analysis was carried out for AAV8 (c, d). Highlighted in orange are peaks derived from empty AAV capsids, blue indicates full capsids. Labels show relative peak area in %.

**Supplementary Figure 2.**
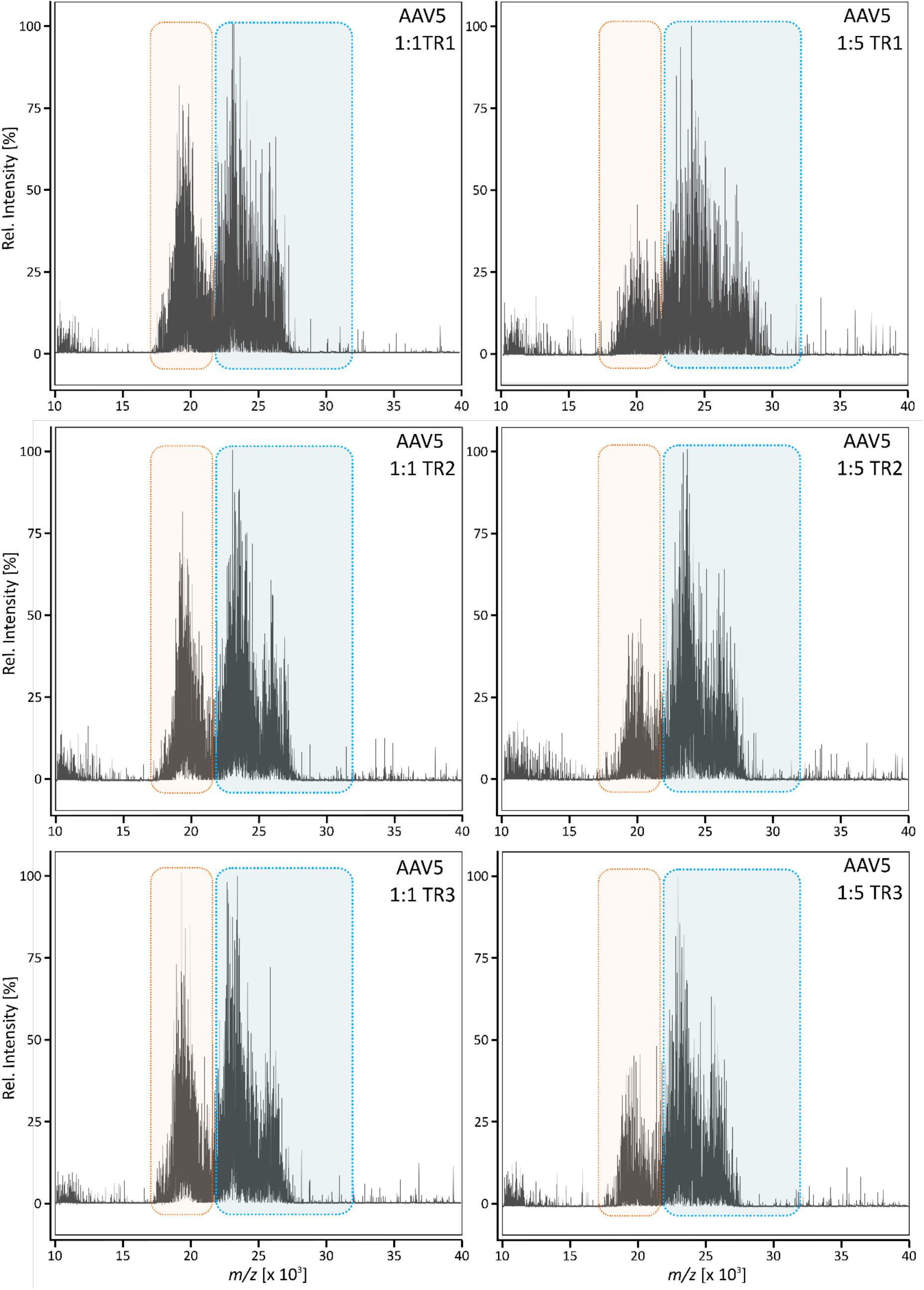
Replicate analysis of full and empty AAV5 capsids using native MS. Signal cluster derived from empty particles are highlighted in orange, full particles are highlighted in blue. Shown are three technical replicates (TR1-3) using a 1:1 or 1:5 mixture of full and empty reference material, respectively.

**Supplementary Table 1.** Empty to full ratio assessment of AAV5. AAV reference material was mixed in a ratio of 1:1 and 1:5, respectively. Samples were analyzed in triplicate via native MS and resulting data was analyzed using ImageJ. Signal clusters corresponding to empty and full capsids were measured and subsequently used to calculate the ratio of empty to full as well as the percentage of full capsids. Shown are average values (n=3) and corresponding standard deviation (SD).

| **AAV5 1:1** | **Average** | **SD** |
| --- | --- | --- |
| % Full (MS) | 62.37 | 0.78 |
| Ratio | 1.66 | 0.06 |
| **AAV5 1:5** |  |  |
| % Full (MS) | 79.26 | 1.04 |
| Ratio | 3.83 | 0.23 |

